# Food environments and obesity – can’t see the fat for the restaurants?

**DOI:** 10.1101/188177

**Authors:** John R Speakman, Mohsen Mazidi

## Abstract

We recently showed that across the mainland USA there is no association between the density of fast food and full service restaurants and the prevalence of obesity. In a recent editorial it was suggested there are 4 problems with our analysis. The suggested problems were the area of analysis may not reflect adequately the exposure to different outlets, using the absolute numbers of restaurants rather than their ratio, using a global model which assumes the same relationship across all sites and finally the potential for residual confounding. In this short note we address all four of these issues and provide some new analysis of the impact of the ratio of restaurant types on obesity prevalence showing there is only a very weak association (r^2^ = 0.006). We conclude that none of the supposed weaknesses in our original analysis are valid.

---

We have recently shown that there is effectively no association between the density of fast-food (FFR) and full service (FSR) restaurants and the prevalence of obesity across the mainland USA (1). We suggest this lack of association calls into question the likely success of attempting to reduce intake from such establishments as a method to curb obesity. Cummins et al (2) in an editorial that accompanied our paper sought to highlight several limitations with our analysis which they claim undermines the central conclusion. Here we provide some clarification regarding these supposed limitations.

The first claimed limitation is the difficulty concerning definition of the environment to which people are exposed. This is obviously an important issue. There is no point, for example, counting the number of FFRs within 1km of a persons home, if they routinely travel 10km from home to eat at FFRs. For the data that we analysed, which concerns the county level data for both obesity prevalence and the restaurant densities, our analysis effectively assumed that individuals in a given county exclusively attend restaurants in the county where they live. Since people consume foods at their home and work locales this assumption could easily be violated if the distance people travel to work routinely takes them outside their home county. We were aware of this potential problem, and tested if this is likely to be an issue by analysing the data in a different way for the state of Georgia. For this state we counted not only the numbers of restaurants in each focal county, but also asked if there was a relationship between obesity levels in the focal counties and the numbers of restaurants in the focal county plus all immediately adjacent counties. Using this broader inclusion criterion didn’t alter the relationship we detected (see Mazidi and Speakman [ref1] Figs S1 to S4). In addition we also analysed data at the State level (Mazidi and Speakman [ref1] fig 5) and the same patterns were also present. Hence the pattern we found was robust to the spatial scale of the analysis, indicating that wrongly defining the zone of exposure was probably not an issue.

The second suggested limitation was our use of the absolute numbers of the restaurants, rather than utilising the ratio of the two or the proportion of establishments of a given type, which have proved more significant in studies of outlet exposure on fruit and vegetable intake (3-5). The reason we used the absolute numbers of restaurants is because there is a clear hypothetical link between the absolute numbers of restaurants per capita of population and obesity, as follows. To remain open restaurants need to sell food. Since food purchased in restaurants is mostly consumed rather than wasted, the more restaurants there are in an area the more restaurant food must be consumed per capita. This greater food consumption per capita might then be causally implicated in over-consumption of food by the population eating it, and hence the absolute density of restaurants may be causally linked to obesity.

We agree that there might be a rationale for why intake of fruit and vegetables could be related to the relative densities of healthy and unhealthy food outlets. That is the presence of a high proportion of unhealthy outlets might distract from the healthy eating options (3, 5). In that situation the different outlet types are presumed to have opposing impacts on the outcome variable, which might justify using their relative abundances. In the analysis we performed, however, this rationale does not appear apt, since the densities of both of the restaurant types we considered were hypothesised to be positively linked to obesity. In fact the relationship between the ratio of the two restaurant types and adjusted obesity prevalence is extremely weak (r^2^ = 0.006), although, due to the large sample size this was significant (F_1, 2944_ =17.3, p < 0.001) (Figure 1). The effect size was also very small: a 55 fold change in the ratio of FFR to FSR was correlated with an average change in adjusted obesity prevalence of just 0.5%.

**Fig. 1.**
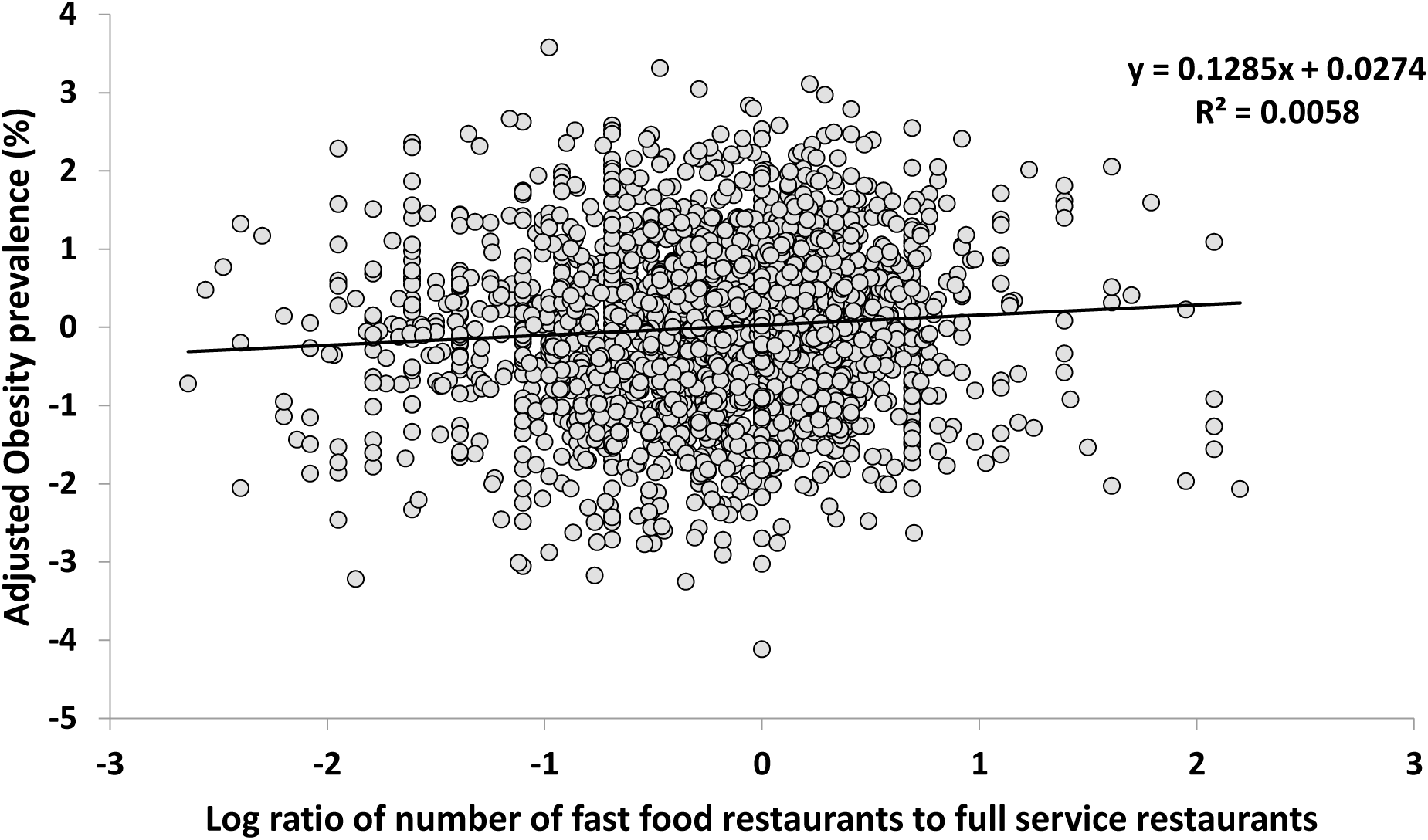
Association between the ratio of fast food to full service restaurants (log transformed) and the prevalence of obesity, adjusted for age, poverty, ethnicity, education, income, insurance status, unemployment and access to recreation facilities (%). Plot shows the county level data across the mainland USA (n = 2945) and the fitted regression.

The third point is that we used a global model to analyse the data which assumes a common relationship exists across the entire area. This is a valid criticism but is easily addressed by asking whether the same absence of a relationship across the entire US is also evident in the individual states. When we performed such an analysis (Mazidi and Speakman: supplementary table 1) we also found no significant association between FFR density and obesity levels in 44 of the 48 states that contributed to the original data. Moreover, we showed in a previous paper using a variogram analysis that there is no spatial autocorrelation in residual obesity levels, indicating the county is an appropriate level for analysis of factors driving obesity relevance (6).

Finally, Cummins et al (2) raise the issue of potential residual confounding. Residual confounding occurs when a relationship is found between two variables A and B, but this is because both A and B are correlated with an unaccounted for confounding variable C. They suggest, for example, that levels of physical activity might be a relevant confounding variable in our study. We agree for FSR the very weak *negative* relationship to obesity prevalence is likely due to some residual confounding, although not necessarily engagement in physical activity. However, for FFR this argument cannot be correct, because for FFR there was no significant association to obesity prevalence. The absence of a relationship between 2 variables cannot be caused by residual confounding of a third variable. The absence of a relationship of obesity prevalence to FFR densities could be because physical activity engagement is a more significant independent factor driving obesity prevalence, hence the much smaller impact of restaurant exposure fails to reach significance. But that was the whole point of our paper. Other factors are far more important for obesity prevalence than exposure to fast food and full service restaurants.

## Abbreviations

FFR: fast food restaurant
FSR: full service restaurant

## Conflict of Interest

None

